# Distinct patterns of surround modulation in V1 and hMT+

**DOI:** 10.1101/817171

**Authors:** Gorkem Er, Zahide Pamir, Huseyin Boyaci

## Abstract

Modulation of a neuron’s responses by the stimuli presented outside of its classical receptive field is ubiquitous in the visual system. This “surround modulation” mechanism is believed to be critical for efficient processing and leads to many well-known perceptual effects. The details of surround modulation, however, are still not fully understood. One of the open questions is related to the differences in surround modulation mechanisms in different cortical areas, and their interactions. Here we study patterns of surround modulation in primary visual cortex (V1) and middle temporal complex (hMT+) utilizing a well-studied effect in motion perception, where human observers’ ability to discriminate the drift direction of a grating improves as its size gets bigger if the grating has a low contrast, and deteriorates if it has a high contrast. We first replicated the findings in the literature with a behavioral experiment using small and large (1.06 and 8.05 degrees of visual angle) drifting gratings with either low (2%) or high (99%) contrast presented at the periphery. Next, using functional MRI, we found that in V1 with increasing size cortical responses increased at both contrast levels, but they increased more at high contrast. Whereas in hMT+ with increasing size cortical responses remained unchanged at high contrast, and increased at low contrast. These findings show that surround modulation in V1 and hMT+ are distinct. Furthermore these findings provide evidence that the size-contrast interaction in motion perception is likely to originate in hMT+.

## 1 Introduction

Visual neurons respond to stimuli only within their classical receptive fields (RF) when these stimuli are presented in isolation. If, however, the RF and its surround are stimulated together, the response patterns of the neurons alter. This surround manipulation is found in many levels of the visual hierarchy (Angelucci et al., 2017). Yet many questions about the mechanism remain open. Most importantly, even though surround modulation is heavily studied in primary visual cortex (V1), it is not clear whether the basic principles of the mechanism remains the same for a variety of stimuli in other visual areas. Here, to tackle this question we used a well-known perceptual effect in motion perception and investigated surround modulation in V1 and human middle temporal complex (hMT+).

In the aforementioned perceptual effect, as the size of a drifting grating increases, discriminating its motion direction becomes harder if it has high contrast, but easier if it has low contrast (Tadin, Lappin, Gilroy, & Blake, 2003). This perceptual effect has been attributed to the surround modulation of neuronal populations in hMT+, which is one of the central cortical areas in motion processing (‘MT-hypothesis’, Tadin et al., 2003). According to the MT-hypothesis, discriminating motion direction of a high-contrast grating becomes harder owing to the suppressive effects of surround stimulation (i.e. surround suppression). For a low contrast grating, on the other hand, motion direction discrimination becomes easier as its size gets larger owing to facilitative effects of surround stimulation (i.e. surround facilitation). The MT-hypothesis has been later supported by functional magnetic resonance imaging (fMRI) findings (Turkozer, Pamir, & Boyaci, 2016; Liu, Haefner, & Pack, 2016; Schallmo et al., 2018). Furthermore, disrupting hMT+ activity with application of TMS resulted in decreased surround suppression for high-contrast stimuli (Tadin, Silvanto, Pascual-Leone, & Battelli, 2011).

Previous studies, however, were not able to address the question of the role of other cortical areas in the observed perceptual effect. For example, neuronal correlates of spatial suppression and facilitation within the hMT+ might be inherited from earlier visual areas, most notably from V1. This possibility has not been systematically analyzed in humans yet. One major cause of this was related to methodological limitations: Previous studies investigating the size-contrast interaction in motion perception used foveally presented stimuli, which activate neurons in the so-called foveal confluence, where borders of V1, V2, and V3 are difficult to draw using fMRI (e.g. Turkozer et al., 2016; Schallmo et al., 2018). Thus, in those studies it was not possible to confidently investigate the activity of V1 along with hMT+. In the current study we overcome this limitation by presenting the stimuli at the periphery, where it becomes straightforward to identify regions of interest in different early visual areas.

We first conducted a behavioral experiment to ensure that the perceptual effect persists when the stimulus is presented at the periphery. After ensuring that the perceptual effect is present, using fMRI we investigated the neuronal responses within hMT+ and V1 in response to drifting gratings in varying contrast and size levels.

## 2 Experiment 1: Behavioral Experiment

### 2.1 Methods

#### 2.1.1 Participants

Eleven participants, including the authors ZP and GE, participated in the experiment (seven female; age range: 19-28). All participants reported normal or corrected-to-normal vision, and had no history of neurological or visual disorders. Prior to the experimental sessions participants gave their written informed consents. Experimental protocols and procedures were approved by the Human Ethics Committee of Bilkent University.

#### 2.1.2 Stimuli, Experimental Procedures, and Analyses

Visual stimuli were presented on a CRT monitor (HP P1230, 22 inch, 1600×1200 resolution, 120 Hz). Participants were seated 75 cm from the monitor, and their heads were stabilized using a chin rest. Responses were collected via a standard computer keyboard. A gray-scale look-up table, prepared after direct measurements (Spectro-CAL, Cambridge Research Systems Ltd., UK), was used to ensure the presentation of correct luminance values. The experimental software was prepared by us using the Java programming platform.

Stimuli were horizontally oriented drifting sine wave gratings (spatial frequency: 1 cycle per degree) weighted by two-dimensional isotropic Gaussian envelopes. Two size- and contrast-matched gratings were simultaneously and briefly presented on a mid-gray background (40.45 cd/m^2^) at +/− 9.06 degrees of horizontal eccentricity (the visual angle between the central fixation and the center of the gratings). Each grating drifted within the Gaussian envelope (starting phase randomized) at a rate of 4 degree/s either upward or downward. Participants reported whether or not the gratings drifted in the same direction, while maintaining fixation at the central fixation mark. After responding, participants received an auditory feedback (auditory tone of 200 ms duration, 300 Hz for correct and 3800 Hz for incorrect answers). Two size levels (small: 1.67, large: 8.05 degrees visual angle in diameter) and two contrast levels (2% and 99% Michelson contrast) were tested (4 experimental conditions in total). Each condition was blocked in a separate session of 160 trials, and the sessions were randomly ordered for each participant. Participants completed a short practice session before beginning an experimental session. Based on their performance in the practice session, initial presentation duration parameter for the experimental sessions were selected per participant. For the ensuing trials, presentation duration was manipulated adaptively with a two interleaved 3-up 1-down staircase procedure. One staircase started from a relatively short duration, the other started from a longer duration. There were 80 trials in each staircase.

Psychometric functions were fit to the data using the Palamedes toolbox (Kingdom & Prins, 2010) in Octave (http://www.octave.org) for each observer and condition. Duration thresholds (79% success rate) and standard errors were estimated. Repeated-measures analysis of variances (ANOVA) with two factors (size and contrast) was performed to compare the thresholds at group level using SPSS Version 19 (SPSS Inc., Chicago, IL). Additionally, to facilitate drawing links between the behavioral results and fMRI findings, we calculated “sensitivity” values, defined as 1/threshold. Next using the sensitivities for large and small Gabors, *S*_*L*_ and *S*_*S*_ respectively, we computed a size index (*SI*) defined as *SI* = *S*_*L*_ − *S*_*S*_. A positive *SI* means increased sensitivity with increasing size (spatial facilitation), a negative *SI* means decreased sensitivity with increasing size (spatial suppression). We compared the *SI* values to “0” by applying one-sample two-tailed Student’s t-test in SPSS. Also, SI values for low and high contrast conditions were compared using two-tailed paired-samples t-test.

### 2.2 Results

We measured duration thresholds for accurately judging the drift direction of Gabor patches presented at the periphery at two size (1.67 and 8.05 degrees) and contrast levels (2% and 99%). Results are shown in Figure 1. Analyses showed that main effect of contrast was statistically significant (F(1,10) = 12.16, *p* < 0.01) and main effect of size was close to significance (F(1,10) = 4.49, *p* = 0.06). Also, the interaction between contrast and size was statistically significant (F(1,10) = 33.96, *p* < 0.001). Bonferroni corrected pairwise comparisons showed that for the high-contrast gratings thresholds increased with size (t(10) = −4.1; *p* < 0.01; small stimuli: M = 39.83, SEM = 2.24; large stimuli: M = 99.8, SEM = 13.02) whereas for the low-contrast gratings the thresholds decreased with size (t(10) = 6.03; *p* < 0.001; small stimuli: M = 107.9, SEM = 6.23; large stimuli: M = 81.18, SEM = 6.22). Consistent with the pairwise comparisons applied to the raw threshold values, one-sample two-tailed Student’s t-tests showed that the size index (*SI*, see methods) was significantly lower than zero for the high-contrast gratings (t(10) = −5.53; *p* < 0.001; M =−0.014, SEM = 0.002) whereas it was significantly higher than zero for the low-contrast gratings (t(10) = 5.56; *p* < 0.001; M =0.003, SEM = 0.0006). Also, two-tailed paired-samples Student’s t-tests showed that SI was significantly higher for low-contrast stimuli compared to that for high-contrast stimuli (t(10) = 6.97; *p* < 0.001). These results clearly replicate the size–contrast interaction in motion perception when stimuli is presented at the periphery.

**Figure 1:**
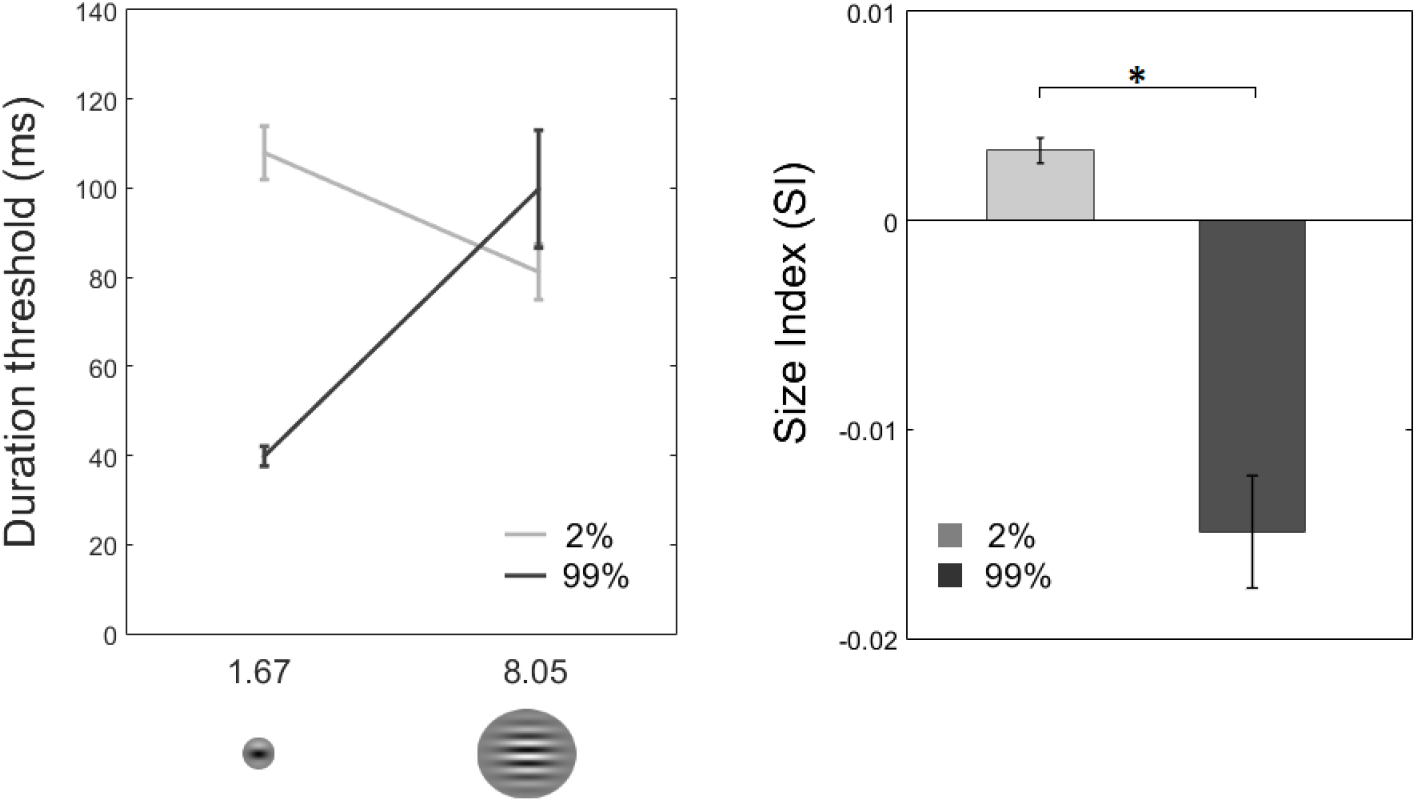
Left plot shows group mean (*N* = 11) of duration thresholds. For low-contrast stimuli, discrimination threshold decreases as size gets bigger. On the contrary, for high-contrast stimuli, discrimination threshold increases as size gets bigger. Right plot shows mean size indices (*SI*s) for 2% and 99% contrast levels. *SI* is defined as the difference in sensitivity (1 / threshold) between large and small Gabor patches. For low-contrast stimuli, *SI* is positive which indicates that sensitivity increases as size gets bigger, i.e. spatial facilitation. On the contrary, for high-contrast stimuli, sensitivity decreases as size gets bigger, i.e. spatial suppression. These results replicate the size–contrast interaction in motion perception when stimuli is presented at the periphery. Error bars represent ±SEM. (\**p* < 0.001).

## 3 Experiment 2: Functional MRI

### 3.1 Materials and Methods

#### 3.1.1 Participants

Six volunteers (age range: 23-26; mean age: 25; three male) participated in the experiment, three of whom also participated in the behavioral experiment. All participants had normal or corrected-to-normal vision and had no history of neurological or visual disorders. Participants gave their written informed consents prior to the fMRI sessions. Experimental protocols and procedures were approved by the Bilkent University Human Ethics Committee.

#### 3.1.2 Data Acquisition & Experimental Setup

MR images were collected in the National Magnetic Resonance Research Center (UM-RAM), Bilkent University on a 3 Tesla Siemens Trio MR scanner (Magnetom Trio, Siemens AG, Erlangen, Germany) with a 32-channel array head coil. MR sessions started with a structural run followed by two region of interest (ROI) localizer and four experimental functional runs, totaling approximately 1 hour in duration. One localizer run was used to identify the hMT+ region, the other one was for localizing the sub-regions of hMT+ and V1 that process the input from the visual field that correspond to the position and size of the small Gabors (see below 3.1.3 “Visual Stimuli & Experimental Design”). Structural data were acquired using a T1-weighted 3-D anatomical sequence (TR: 2600 ms, spatial resolution: 1 mm^3^ isotropic, number of slices: 176). Functional images were acquired with a T2*-weighted gradient-recalled echo-planar imaging (EPI) sequence (TR: 2000 ms; TE: 35 ms; spatial resolution: 3×3×3 mm^3^; number of slices: 30; slice orientation: parallel to calcarine sulcus). Visual stimuli were presented on an MR-safe LCD Monitor (TELEMED PMEco, Istanbul, Turkey; 32 inch; resolution: 1920×1080; vertical refresh: 60 Hz). The monitor was placed near the rear end of the scanner bore, and viewed by the participants from a distance of 165 cm via a mirror mounted on the head coil. The stimuli were generated and presented using Python and the Psychopy package (Peirce, 2009).

#### 3.1.3 Visual Stimuli & Experimental Design

Visual stimuli were drifting Gabor patches as in the behavioral experiment. Two size (small: 1.67 degree, large: 8.05 degree) and two contrast levels (2% and 99%) were tested. Due to the limits of the visual display system, gratings were presented at +/− 8.02 degrees of horizontal eccentricity (was 9.06 degrees in the behavioral experiment), and drifted with a rate of 6 degree/s (was 4 degree/s in the behavioral experiment) either upward or downward for the duration of 12 seconds. Both Gabor gratings drifted in the same direction simultaneously, and alternated direction every two seconds to avoid motion adaptation.

A functional run was composed of “active” and “control” blocks, each lasting for 12 seconds. In the active blocks, drifting Gabor patches and a central fixation mark were presented, whilst in the control blocks, only the fixation mark remained visible. In alternating active blocks, small and large drifting Gabor patches were shown, each repeated for 6 times in a run. Contrast level was kept constant within a run. Two experimental runs were conducted for each contrast level in a session. The runs started with an initial blank period of 24 seconds to allow hemodynamic response to reach a steady state. The total duration of a functional run was around 5 minutes. Figure 2 shows the schematic representation of an experimental run.

**Figure 2:**
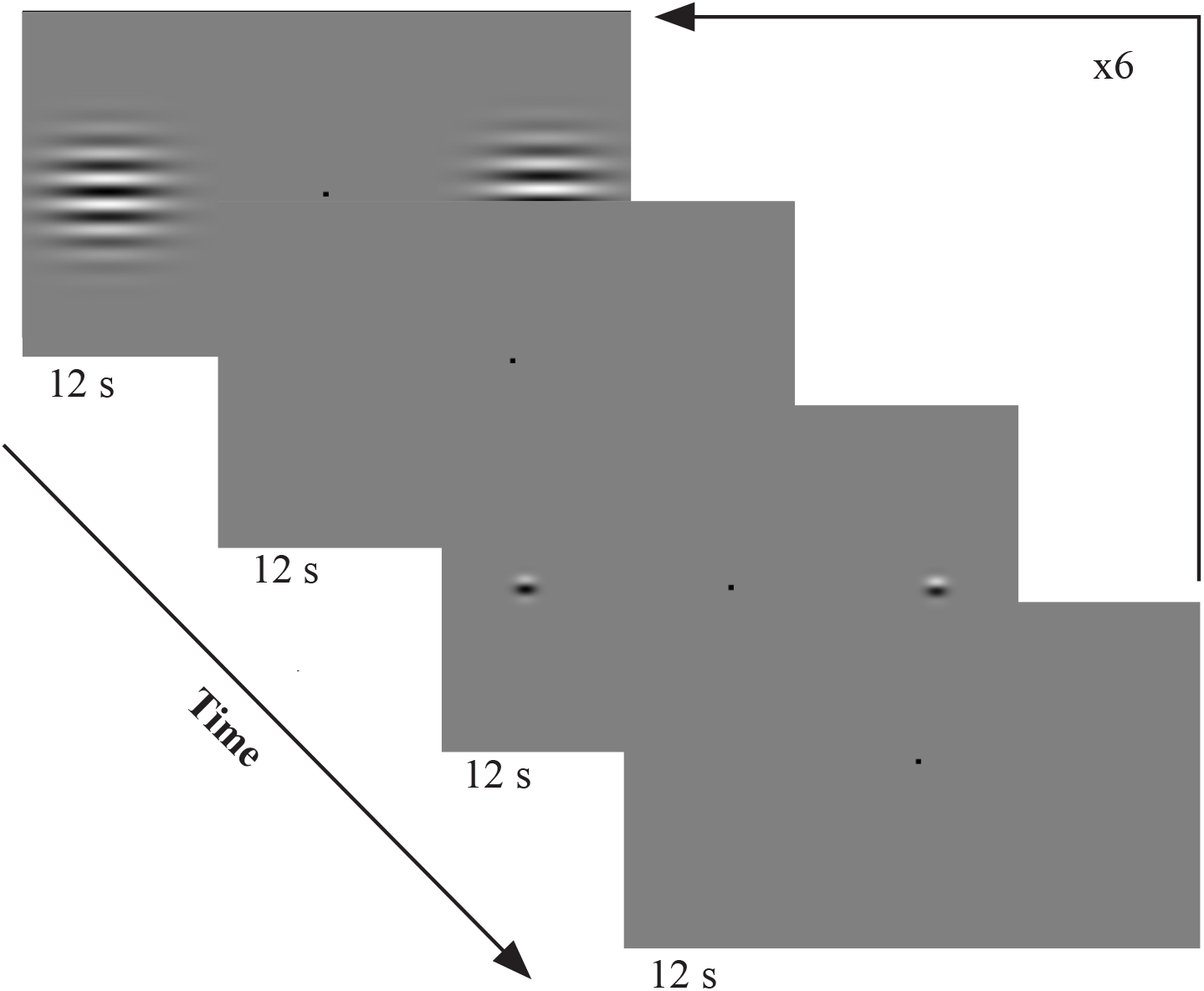
Schematic representation of the visual paradigm of a single cycle in the fMRI experiment. This cycle is repeated for 6 times within a run. Large (8.05 degree) and small (1.67 degree) drifting Gabor patches were presented in alternating active blocks. The contrast was kept constant in a run (2% or 99%), and two runs were conducted for each contrast. Participants were required to keep fixation at the central mark, and perform a demanding fixation task.

Both to ensure fixation and to control for spatial attention, participants were asked to perform a demanding fixation task throughout an entire functional run (all participants achieved a mean accuracy rate of over 90%, which was the predetermined threshold to discard the participant’s data). The color of the fixation mark (0.3-degree solid square) changed randomly from its original color (gray) to either red or yellow for the duration of 50 ms at the randomly designated interval between 250 to 1500 ms. The participants’ task was to report the changes in the color of the fixation mark by pressing the designated button on an MR-safe response button-box (Fiber Optic Response Devices Package 904, Current Designs).

#### 3.1.4 hMT+ Identification

We identified the hMT+ complex in a separate run using the established methods in literature (Huk, Dougherty, & Heeger, 2002; Dukelow et al., 2001; Smith, Wall, Williams, & Singh, 2006). Specifically, we presented the participants fields of moving dots while acquiring functional MR images. The dot fields were comprised of 100 dots on a black background within an 8-degree diameter circular aperture. The centers of the fields were 8.02 degrees to the left and right of the fixation point. The dots moved in three different trajectories: radial (expanding - contracting), cardinal (left-right; up-down), and angular (clockwise-counterclockwise). Motion direction changed every two seconds to prevent adaptation. BOLD responses were collected for three types of configurations, each presented for 12-seconds: right field dynamic (left static), left field dynamic (right static), and both fields static. This cycle of the presentation was repeated eight times in a run. We used general linear model (GLM) to contrast the BOLD responses during dynamic and static presentations. Voxels that respond more strongly to contralateral dynamic compared to static stimuli at the ascending tip of the inferior temporal sulcus were identified as hMT+.

#### 3.1.5 ROI Localization within hMT+ and V1

Within hMT+ and V1, subregions which are selectively more responsive to the visual field that correspond to the locations of the small Gabors in the experiment were identified using another localizer run. This localizer run was composed of 12s active and rest blocks. In the active blocks participants saw drifting Gabor patches whose size, position and speed matched the small stimuli in the experimental runs. The contrast of the Gabor patches, however, was 60%. Throughout the entire run participants were required to maintain central fixation and perform a demanding fixation task as described before. All subsequent analyses were performed on the experimental data extracted from these ROIs.

##### ROI within hMT+

We first created masks using the hMT+ regions functionally identified as explained before. We then localized the hMT+ ROIs within these masked regions. Specifically we identified the voxels that are selectively more responsive to the drifting Gabor patch by contrasting the responses between active and control blocks using GLM.

##### ROI within V1

We identified V1 ROI using the data from the localizer run with the aid of anatomical landmarks. Specifically, using GLM, cortical regions responding preferentially more in the active blocks when contrasted with control blocks and located anteriorly in the calcarine sulcus were identified as the V1 ROI.

#### 3.1.6 Analyses

Anatomical and functional data were preprocessed and analyzed using the BrainVoyager QX software (Brain Innovation, The Netherlands). Preprocessing steps for the functional images included head motion correction, high-pass temporal filtering and slice scan time correction. T1-weighted structural images were transformed into the AC-PC plane, and aligned with the functional images. For each brain, the border between white matter and cortex was drawn, and an inflated three-dimensional model of the cortex was generated. Functional maps were projected onto the inflated cortex to aid the visualization of subsequent analyses. hMT+, and subregions within hMT+ and V1 were identified with GLM as described before using BrainVoyager.

Statistical analyses were performed on BOLD responses computed by using the beta weights calculated with GLM within the ROIs. To further quantify the changes in BOLD response evoked by an increase in stimulus size, and to draw links with behavioral results, we calculated an fMRI size index (*SI*) defined as *SI* = *B*_*L*_ − *B*_*S*_, where *B*_*L*_ and *B*_*S*_ are the BOLD responses for large and small gratings, respectively. A positive SI denotes an increased BOLD response with increase in size (surround facilitation), and a negative SI denotes decreased BOLD response (surround suppression). To compare the BOLD responses at the group level, a 2 × 2 repeated-measures ANOVA was conducted with contrast level (low and high) and size of the stimuli (small and large) as factors. We applied one-sample two-tailed Student’s t-test to compare *SI*s to zero at each contrast level. We also performed paired samples t-test to compare the *SI*s to each other (low versus high contrast). Statistical analyses were conducted using JASP Version 0.8.5 (JASP Team, 2018).

### 3.2 Results

In this experiment, we recorded and analyzed BOLD responses in hMT+ and V1 while the observers viewed peripherally presented drifting Gabor patches. We compared the magnitudes of BOLD responses between small and large Gabors at two contrast levels within predefined ROIs that correspond to the location and size of small stimuli (i.e. “center”). Because our goal here is to measure the modulatory effect of surround stimulation on the responses of the neurons whose classical RF centers are inside the visual space that correspond to the small Gabor patch. BOLD response differences evoked by presenting the large and small-sized stimuli would highlight the suppressive or facilitative influence of the surround on the center.

#### 3.2.1 hMT+

Figure 3 (left plot) shows the results from hMT+ ROI. Increasing the size of the stimuli resulted in increased BOLD response, when stimuli had low contrast. Conversely, increasing size of the stimuli resulted in no change in BOLD response when stimuli had high contrast. To test whether stimulus contrast and size affect the magnitude of BOLD response significantly, we applied a 2 × 2 repeated-measures ANOVA for the contrast (2% and 99%) and size (1.67 and 9.05 degree) as factors. Results revealed a no main effect of contrast (F(1,5) = 3.69, *p* = 0.113), while we found a main effect of size (F(1,5) = 19.41, *p* = 0.007), as well as an interaction between size and contrast (F(1,5) = 10.257, *p* = 0.024) at hMT+.

**Figure 3:**
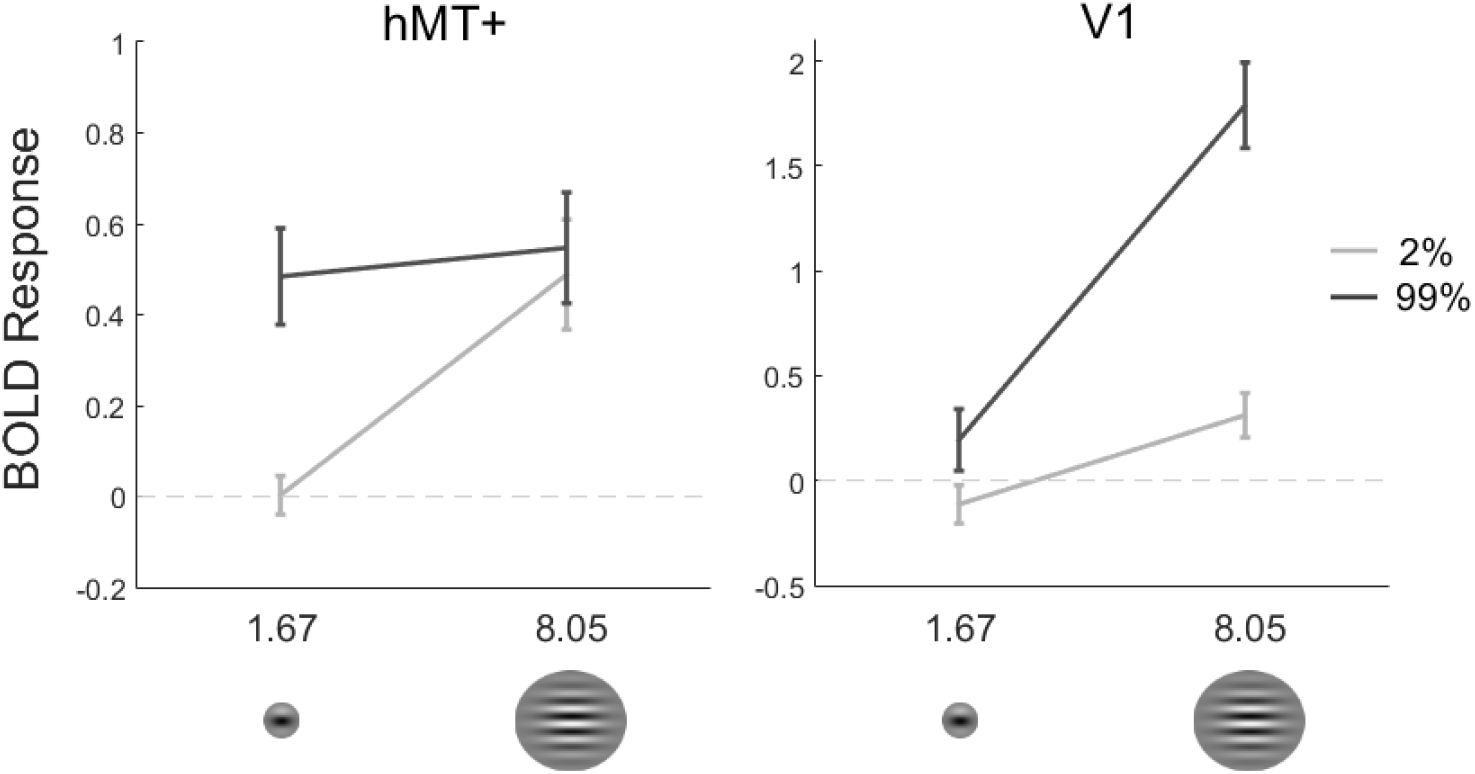
Group mean (*N* = 6) of BOLD responses from hMT+ and V1. BOLD responses to small and large stimuli with high and low contrast were extracted from ROIs that correspond to the location and size of the small stimuli (identified with an independent functional localizer run). Average responses from the ROIs were computed for each subject, then the group mean was calculated. Error bars represent SEM.

To further investigate the patterns of results, we computed size indices (*SI*s), defined as the difference between the BOLD responses to large and small Gabors (see Methods). Figure 4 shows individual *SI* values for all participants, as well as the mean *SI*. We found that *SI* was significantly different (greater) than zero at low contrast (*M*_*SI*_ = 0.486, SEM = 0.110; one-sample t-test, t(5) = 4.42, *p* = 0.007). On the other hand, the *SI* at high contrast was not significantly different than zero (*M*_*SI*_ = 0.063, SEM = 0.066; one-sample t-test, t(5) = 0.95, *p* = 0.385). Furthermore, based on the results in literature (Turkozer et al., 2016; Schallmo et al., 2018), we expected a larger *SI* for low contrast compared to high contrast. Indeed, paired sample t-test results revealed that the *SI*s were statistically significantly different, the *SI* for low-contrast being greater than that for the high-contrast (t(5) = 3.20, *p* = 0.024).

**Figure 4:**
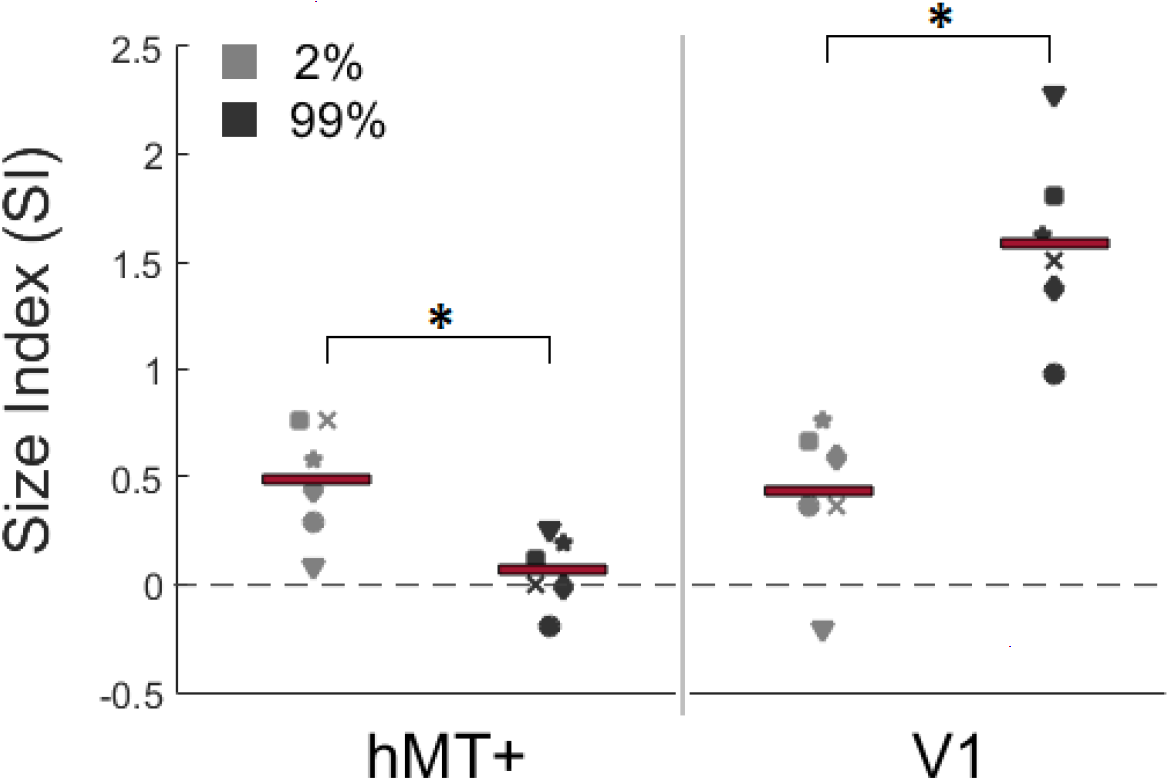
fMRI size indices (*SI*) in hMT+ and V1 for individual participants, and the group mean (red bars). *SI* is defined as the difference in BOLD response between large and small stimuli. A positive *SI* indicates surround facilitation, a negative one surround suppression. Compare to behavioral results in Figure 1, right plot. (\**p* < 0.05).

#### 3.2.2 V1

We next analyzed how the size and contrast of stimuli affect BOLD responses within the V1 ROI. Figure 3 (right plot) shows the responses for each condition. Results showed that BOLD responses increased significantly with size both at high and low contrast conditions. Critically, this increase was greater when the stimuli had high contrast compared to low contrast. This pattern was inconsistent with the perceptual effect, and surprisingly it was different than the pattern observed in hMT+. We applied two-way repeated-measures ANOVA with the contrast (low and high) and size (small and large) as factors to investigate the role of contrast and size on the BOLD response. ANOVA revealed a main effect of contrast, (F(1,5) = 17.83, *p* = 0.008), and size (F(1,5) = 148.94, *p* < 0.001), as well as the interaction between size and contrast (F(1,5) = 17.96, *p* = 0.008).

As we did for the hMT+ data, here too we performed further analyses on *SI*s. Figure 4 shows *SI*s plotted for individual participants, as well as the group mean. At low contrast, group mean of *SI* was significantly greater than zero (*M*_*SI*_ = 0.425, SEM = 0.143; one sample t-test, t(5)= 2.98, p = 0.031). Average *SI* value was positive at high contrast, as well (*M*_*SI*_ = 1.596, SEM = 0.178; one sample t-test, t(5) = 8.98, *p* < 0.001). Furthermore, we performed paired-sample t-test, and found that the *SI* value for low contrast was significantly lower than the *SI* value for high contrast (t(5) = −4.24, *p* = 0.004).

### 3.3 Linking Behavioral and fMRI Results

To draw a link between the behavioral and fMRI results, we first assume that the amount of neuronal responses can be approximated by a monotonically increasing function of sensitivity (1/duration threshold). Further, we assume a linear relation between neuronal activity and the BOLD fMRI response (see e.g. Boynton, Demb, Glover, & Heeger, 1999). Thus, if an area is involved in the processes related to the perceptual effect, we expect an increase in fMRI BOLD response in that area as the behavioral sensitivity gets better. Comparing the duration thresholds (Figure 1, left plot), and BOLD responses (Figure 3), we see that the hMT+ activity captures the behavioral results for low contrast stimulus; for high contrast stimulus, however, there seems to be a slight disagreement (see Discussion for possible reasons for this). The V1 activity, on the other hand, is completely inconsistent with the behavioral results.

This pattern becomes more clear when the size indexes are compared (Figure 1 right plot, and Figure 4). Overall the response patterns in hMT+, but not in V1, agree well with the behavioral results.

## 4 Discussion

We demonstrated that surround modulation found in hMT+ complex, but not in V1, agrees with the size-contrast interaction in motion perception. First in a behavioral experiment we measured the temporal thresholds for successfully detecting the direction of motion of drifting Gabor patches presented at the periphery. We found that the thresholds decreased with size (i.e. increased sensitivity, spatial facilitation) if the target has low contrast (2%). Conversely, we found that the thresholds increased with size (i.e. decreased sensitivity, spatial suppression) if the target has high contrast (99%). These results were in good agreement with literature (e.g., Tadin et al., 2011). Next, we recorded BOLD fMRI responses while participants viewed high- and low-contrast, small and large drifting Gabor patches presented at the periphery. In hMT+ we found that BOLD responses significantly increased with size for the low contrast Gabor patches (surround facilitation), and remained unchanged for the high contrast Gabor patches. In V1, however, BOLD responses increased for both high and low contrast Gabor patches; and critically the increase was stronger for the high contrast stimuli. Overall, we contend that the activity patterns in hMT+, but not in V1, reflect the perceptual effect.

The BOLD responses we found in hMT+ for high-contrast stimuli may at first seem inconsistent with the behavioral results. Specifically, behaviorally the sensitivity decreases with size, whereas the BOLD responses in hMT+ remains unchanged instead of also decreasing. We believe that there may be several reasons for this. Firstly, because we were interested in the modulatory effects of the surround on the neurons that were processing the center, we sought to identify regions of cortex that process the part of the visual field that correspond to the position and size of the small stimuli (center). To do this, we identified the voxels that responded more strongly to small drifting Gabors compared to a blank screen. Given the large number of neurons in an fMRI voxel (about a million), and their possible heterogeneity, it is likely that neurons with larger receptive fields that respond directly to both small (center) and large (center + surround) stimuli could have been included in our ROI. This scenario is especially likely in hMT+ where RF sizes are usually much larger. Those neurons could have responded more strongly during the presentation of the large stimulus, hiding the effect of response reduction in the neurons that respond only to the center. Alternatively, the observed discrepancy could be because of saturation of BOLD responses at high contrasts (Tootell et al., 1995; Buracas & Boynton, 2007), or because of the commonly observed non-linear relations between the stimulus energy and BOLD responses (Logothetis & Wandell, 2004). Considering these possibilities, we believe that assessing the neuronal responses using the Size Index (*SI*) is more appropriate, because *SI* better represents the overall pattern of interaction between size and contrast. Based on the *SI*s, we see that hMT+ activity agree with the perceptual sensitivity. The pattern in V1, however, is completely at odds with the perceptual effect.

Surround modulation in hMT+ has been previously claimed to underlie the size-contrast interaction in motion perception (Tadin et al., 2003). Two recent neuroimaging results landed support for this hypothesis (Turkozer et al., 2016; Schallmo et al., 2018). Specifically, in both studies, the response patterns in hMT+ were found to reflect the size-contrast interaction. The neuronal activity in hMT+, however, could be inherited from earlier areas, particularly the primary visual cortex (V1). To examine this possibility required investigating the responses in earlier areas. This could not be done previously with fMRI due to methodological limitations. In both studies stimuli were presented at the fovea (Turkozer et al., 2016; Schallmo et al., 2018). This part of the visual field is mapped onto the so-called foveal confluence at the occipital pole, where it is difficult to reliably distinguish V1, V2, and V3 using the standard retinotopic mapping techniques (e.g. Engel, Glover, & Wandell, 1997). Therefore neither of the previous studies could argue strongly that the size-contrast interaction in motion perception did not originate in earlier visual areas (Turkozer et al., 2016; Schallmo et al., 2018). In the present study we have successfully avoided this limitation by presenting the stimuli at the periphery, which allowed us to confidently localize ROIs in V1, as well as hMT+.

Our results in hMT+ agree with the results of the two studies introduced above (Turkozer et al., 2016; Schallmo et al., 2018). Turkozer et al. (2016) did not report results from other visual areas, but Schallmo et al. (2018) found surround suppression for all contrast levels in early visual cortex (EVC), which is defined as the sum of V1, V2 and V3 in the foveal confluence. This seems to stand in contradiction to our findings. The major difference between our study and Schallmo et al. (2018) was the position of the stimuli (periphery vs. fovea). Owing to the methodological limitations described before, Schallmo et al. (2018) could report only the aggregated activity of V1, V2, and V3 (EVC). Reporting the averaged activity from EVC, however, may have obscured the facilitation in V1. This is probable, because suppression has been shown to be progressively stronger in V2 and V3 than in V1 (Zenger-Landolt & Heeger, 2003). Alternatively, the difference in surround modulation at the fovea and periphery in V1 could have caused the differences between our results and Schallmo et al. (2018) results (Xing & Heeger, 2000). This will be elaborated further below.

In their study, Schallmo et al. (2018) argue that a single computational mechanism, namely divisive normalization (Heeger, 1992; Reynolds & Heeger, 2009; Carandini & Heeger, 2012), can successfully account for both surround facilitation and suppression (also see Schallmo et al., 2019). Borrowing this idea, we have also tested how well the divisive normalization model could explain our results with the estimated RF center and the surround size at the periphery. We found good agreement between the model and hMT+ responses. But the model could not predict the responses in V1. This could be because of the complex role of eccentricity in mediating surround modulation in V1. For example, divisive normalization may be a good choice to formulate the effects of feedforward mechanisms, but surround modulation, particularly at the periphery, may involve more than these feedforward connections, and include feedback and horizontal mechanisms, as well (Nurminen & Angelucci, 2014; Nurminen, Merlin, Bijanzadeh, Federer, & Angelucci, 2018). There is little doubt that as part of a vastly interconnected network V1 receives feedback from other visual areas (Shao & Burkhalter, 1996), including those for motion processing (Hupé et al., 1998; Ponce, Lomber, & Born, 2008; Paffen, van der Smagt, te Pas, & Verstraten, 2005). Such top-down influences may need to be factored in a computational model for a fuller understanding of surround modulation in V1. Moreover, such a model should also be able to incorporate the effects of attention, which is shown to interact with eccentricity in surround modulation in V1 (Reynolds & Heeger, 2009).

It is worth noting that there seems to be a disagreement between the surround modulation in V1 found using cell-recording methods on animal models and fMRI on humans. Using cell-recording techniques, surround suppression has been routinely shown for high-contrast stimuli in V1 (e.g. Jones, Grieve, Wang, & Sillito, 2001; Angelucci & Shushruth, 2013; Angelucci et al., 2017). Similar to our findings here, Press, Brewer, Dougherty, Wade, and Wandell (2001) reported only facilitation in V1 using flickering checkerboard patterns (but see Zenger-Landolt & Heeger, 2003). These inconsistencies may indicate a species difference, or differences between the methods (i.e. fMRI vs. cell recording, and differences in experimental procedures). Reconciling these differences would require careful and systematic comparison of results using the same experimental conditions.

## 5 Conclusion

Our results provide further evidence that size-contrast interaction in motion perception likely originates in hMT+. In a broader context our results show that surround modulation can be distinct in different components of the biological visual system. The good agreement of hMT+ response patterns in periphery (this study) and fovea (Turkozer et al., 2016; Schallmo et al., 2018) suggests that surround modulation in hMT+ is relatively stable across eccentricity. But this may not be true for the V1 neurons. Overall, these results show that V1 and hMT+ may exhibit different characteristics of surround modulation.

## 6 Author Contributions

ZP and HB conceived the original idea. ZP designed, implemented and conducted the behavioral experiments. ZP and GE designed; GE implemented and conducted the fMRI experiments. GE, ZP, and HB wrote the manuscript.

## 8 Acknowledgments

We thank Michael-Paul Schallmo and his colleagues for letting us use their code for testing the divisive normalization model. We thank Halide Bilge Turkozer for her contribution to designing the experimental paradigm. Author ZP was supported by TÜBİTAK (National Scholarship Program for PhD Students, scholarship id: 2211-E).

